# Fine-tuning TrailMap: The utility of transfer learning to improve the performance of deep learning in axon segmentation of light-sheet microscopy images

**DOI:** 10.1101/2023.10.23.563546

**Authors:** Marjolein Oostrom, Michael A. Muniak, Rogene M. Eichler West, Sarah Akers, Paritosh Pande, Moses Obiri, Wei Wang, Kasey Bowyer, Zhuhao Wu, Lisa M. Bramer, Tianyi Mao, Bobbie Jo Webb-Robertson

**Author notes:** Correspondence should be addressed to T.M. or B.M.W.

## Abstract

Light-sheet microscopy has made possible the 3D imaging of both fixed and live biological tissue, with samples as large as the entire mouse brain. However, segmentation and quantification of that data remains a time-consuming manual undertaking. Machine learning methods promise the possibility of automating this process. This study seeks to advance the performance of prior models through optimizing transfer learning. We fine-tuned the existing TrailMap model using expert-labeled data from noradrenergic axonal structures in the mouse brain. By fine-tuning the final two layers of the neural network at a lower learning rate of the TrailMap model, we demonstrate an improved recall and an occasionally improved adjusted F1- score within our test dataset over using the originally trained TrailMap model.

Availability and implementation: The software and data are freely available at https://github.com/pnnl/brain_ohsu and https://data.pnl.gov/group/204/nodes/dataset/35673, respectively.

## Introduction

Understanding how functional brain states are achieved and how they are modulated by experience, genetics, and pharmacological agents require a reference map of the structural connections between neurons. Such a map will generate insights into the information processing mechanisms of healthy brains but also facilitate research into brain disorders. Such is the goal of the Mouse and Human Connectome Projects [1-3].

Light-sheet fluorescence microscopy (LSFM) is an important tool in this discovery process. Based on wide-field fluorescence technologies, LSFM enables direct visualization of cellular structure on a scale of a few micrometers. LSFM introduced the use of intrinsic optical sectioning capabilities by automating the movement of a sample through a single plane sheet of light. The resulting fluorescence is detected by an orthogonal-placed camera. By only illuminating a thin section of tissue with each pass, photobleaching of the tissue is minimized, yielding a more intense and complete fluorescence signal. Another important parallel advancement is optical tissue clearing. Through the use of organic solvents, lipid removal, or immersion in refractive index matching solutions, brain tissue is rendered effectively transparent, allowing greater contrast of the labeled fluorescent structures [4-6]

Combining LSFM and brain clearing enables the tracing and analyzing of whole brain connectivity at mesoscopic level. However, technical challenges remain to make this high-resolution data meaningful for connectome research, specifically: to stitch together aligned serial sections to construct a 3D image volume; to efficiently annotate voxels of interest (for example, labeling the content as belonging to the axon of a particular neuron); and, registering landmarks in an image volume with a reference atlas so that data from multiple studies might be compared in the same coordinate space [7].

Manual annotation is one approach to addressing some of these challenges. However, annotating a volume of images is a laborious and time-intensive process that can be subjective due to fatigue-related errors and inhomogeneities in the intensity of stained structures resulting in significant variability between experimenters. In addition, the number of image slices that yield a complete volume range from several dozen to over ten thousand, depending on the microscopy system, species, and orientation of the sectioning method [8, 9]. A complete dataset of high-resolution data for a whole brain can be on the order of a terabyte, limiting analysis algorithms to those that do not require the full volume of data to be loaded into run-time memory [10].

Computational methods to automatically stitch, segment, annotate, and register meaningful information in images have made enormous strides in the past decade. Early methods tended to rely on thresholding, clustering, edge detection, region growing, or curve propagation [11, 12]. Algorithms have also been developed to trace axons, such as the all-path-pruning algorithm, which uses existing information such as the starting and end points of neurite structures, to trace 3D axons structures [13]. A significant break-through came in 2015 with the development of a class of convolutional networks (CNNs) that use symmetric contractive and expansive passes over augmented data samples. Such an approach allowed for a balance between localization and context, as well as a robustness to deformations and variance within the examples from which the network learned. The first such class of these networks, U-Net, won the ISBI (IEEE International Symposium on Biomedical Imaging) cell tracking challenge in 2015 by a wide margin [14]. This success yielded a plethora of new U-Net architectures and approaches intended to refine and build on this advancement, but also to address some of the limitations. The first limitation is that CNNs require that their hyperparameters be tuned to obtain optimal performance, which can be considerably time consuming. In addition, the problem of having a sufficient number of hand-annotated datasets with which to train the CNN remains an issue.

TrailMap (Tissue Registration and Automated Identification of Light-sheet Microscope Acquired Projectomes) is a Python package developed for training a a 3D U-net model for axon segmentation by Friedmann et al. [15]. The original TrailMap model was trained with volumes of brain substacks containing labeled serotonergic axons and subsections containing imaging artifacts and background. Each volume contained 3 - 10 labeled slices (Figure 1 shows an equivalent example from our dataset). The package’s evaluation accounts for these sparsely labeled images by masking unlabeled slices. TrailMap generates a one-pixel boundary representing the edges of axons from the images and devalues their misclassification in calculating loss compared to the misclassification of axons. This ensures that a false positive within one pixel of the axon is not strongly penalized, thereby minimizing the effect of pixel-size differences in expert labeling.

**Figure 1.**
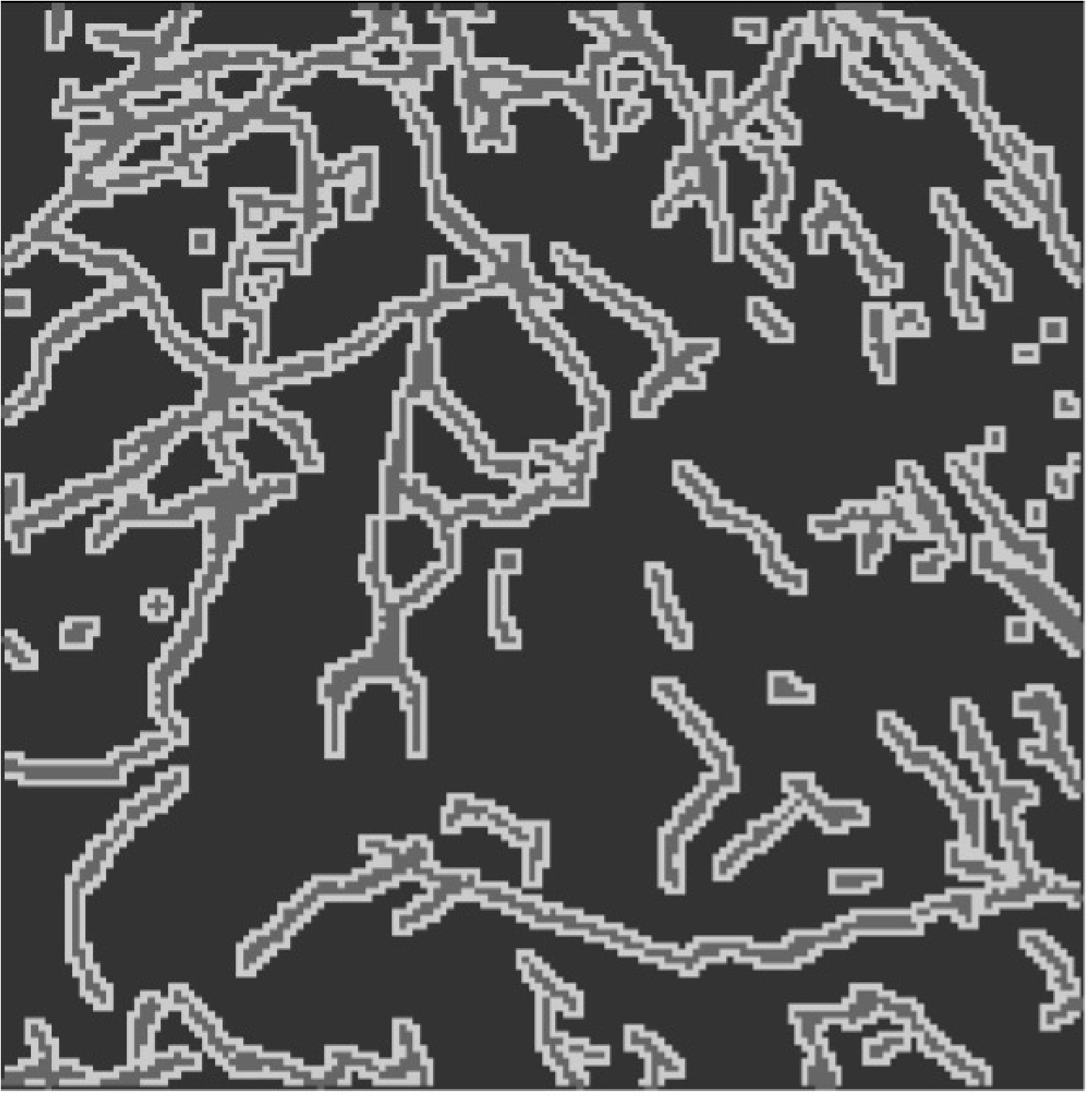
Data Label: An expert-labeled slice where darker regions are background, lighter regions are axons, and the lightest regions are edges.

Technologies such as TrailMap contribute to the growing effort to develop templates for integrating data across multiple non-disruptive 3-dimensional image modalities, combining insights from light-sheet microscopy and MRI into a common coordinate framework [15, 16]. Such co-registration is essential for studying disease progression, developing effective therapies, and translating findings into clinical procedures, for example, delivering brain site-specific drugs and implanting electrodes precisely for recording or stimulation.

Transfer learning can be a useful method in which the features learned from another domain with a richly annotated dataset are used to segment a new smaller dataset outside of the original domain [17]. Fine-tuning the TrailMap model provides an opportunity to exploit the benefits of transfer learning specifically from other images within this domain [15], such as our dataset of labeled brain-wide noradrenergic projections. Indeed, TrailMap includes code and instructions for transfer learning, and the authors already had successfully applied transfer learning to prefrontal cortical projections with 17 training cubes. Our training dataset, however, is much smaller.

We wished to keep the innovative features of TrailMap (e.g., sparse labeling, edges generation, network weights from pretraining), while improving the performance of the fine-tuned model on our dataset. While naturally maintaining the same TrailMap voxel label structure of axons and background, our annotated data involves a different neurotransmitter system (noradrenergic rather than serotonergic), an updated clearing technique, and a different light sheet microscope (LifeCanvas SmartSPIM rather than LaVision Ultramicroscope II light-sheet). Neurons have many diverse morphologies and projections [18], so one implication of using neurons from a different neurotransmitter system is that the original model could improve its performance within the new neurotransmitter system, which might have different axonal projection types, by being fine-tuned.

In optimizing fine-tuning, we incorporated several principles from nnU-net (no-new-U-Net) as modifications in the TrailMap code [19]. nnU-net is a codebase that decides the optimal training values for some parameters based on the characteristics of a database and has yet others that remain fixed regardless of the dataset. While nnU-net was originally developed to optimize training, rather than the fine-tuning of transfer learning, many of these fixed parameters seemed relevant to fine-tuning, including parameters for data augmentation, data foreground sampling, scheduled learning rate, and the inference overlap method. We will refer to modifications inspired by the nnU-net codebase as the “nnU-net modifications” in this paper.

One optimization approach to avoid overfitting is data augmentation. Data augmentation is the process of increasing the amount of training data by generating new instances from existing data; the strategies employed reduce the tendency of a model to overfit. Common techniques include a combination of spatial transformations such as scaling, rotation, reflection, and cropping. TrailMap includes algorithms for rotations and horizontal, vertical, and depth-wise flipping of volumes. We also tested algorithms for elastic deformation as we thought the natural variation in axons might mimic such a transformation. Elastic deformation was not mentioned as an augmentation method in the nnU-net paper, but it is included in the nnU-net codebase [19] and natural variation in axons may also mimic elastic deformations. We did not implement all the augmentation techniques included in nnU-net, including notably, augmentations affecting the intensity of the image by altering the z-score; TrailMap had already determined that z-score normalization, by removing the raw intensity values from the images, reduced the ability of the model to distinguish background from axons.

An illustration of the spatial transformations on an input volume and labels can be seen in Figure 2. It includes an artificial line to illustrate that the volume inputs and labels remain correctly mapped to each other after the transformation.

**Figure 2.**
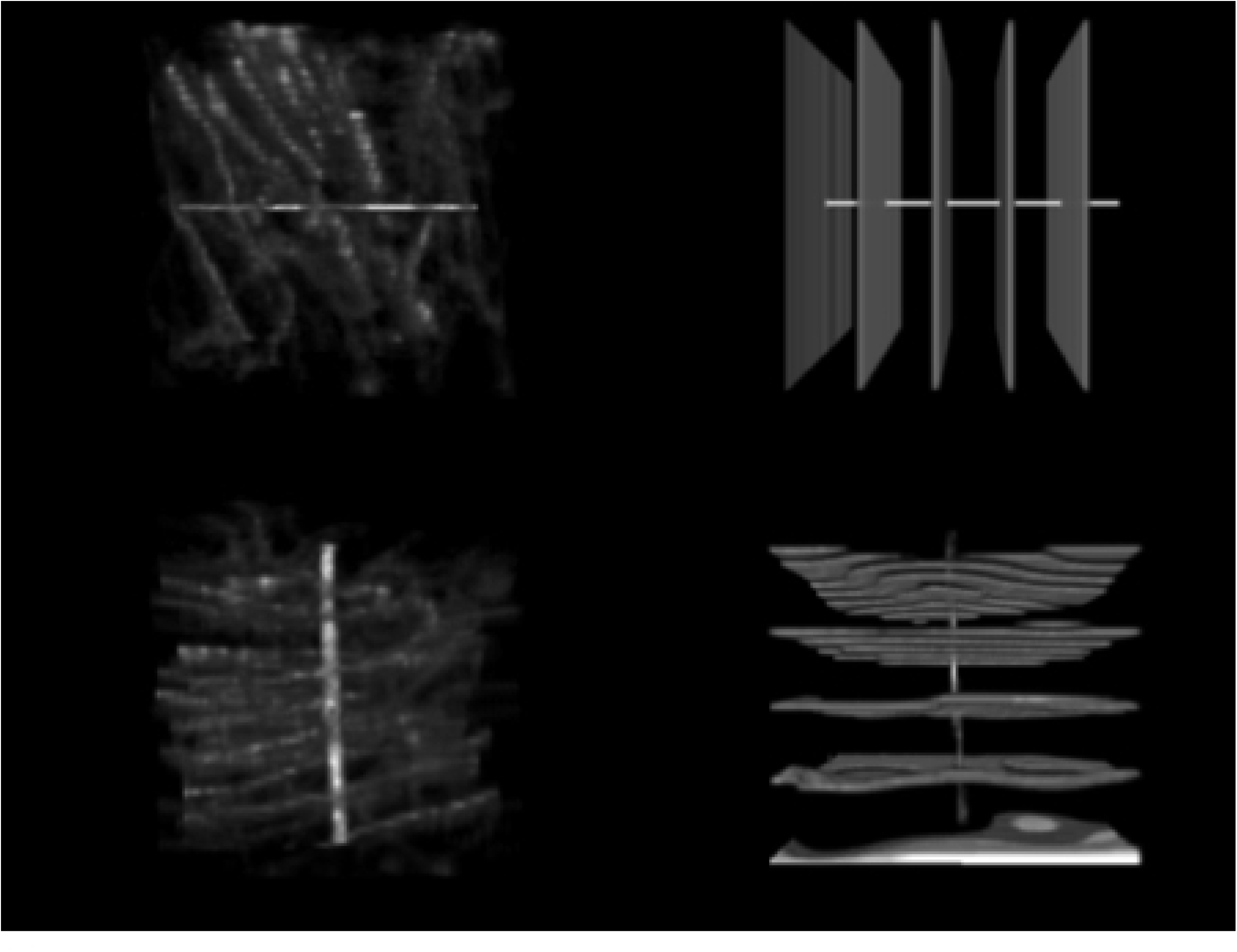
Data Augmentation: A: image inputs (top row) with a horizontal line added for transformation comparison. B: A labeled volume with all labels (axons, edges, and background) in white. C. input transformation D. label transformation. For A and C the intensity is adjusted so 0 as the minimum and 5140 as the maximum (visualized using intensities divided by 65535 and multiplying by 255 within the code, and then adjusted using 0 and 20 as the max and min within Fiji). All images are 18.66 um on the original rotation x and y axis and 20 um on the original rotation z axis and displayed with Fiji. Using 100 pixel^3^ cubes for illustration.

An additional nnU-net modification is to over-sample the 3D volumes from areas in the full training image with foreground features [19]. The foreground features for our study are the axons, which are indicated in the labeled data, so we included over-sampling as well, which we labeled as a nnU-net modification.

nnU-net uses a learning rate that decays during training. We added in an adaptive learning rate that decreases when there is a prolonged period of no improvement to the validation loss, which we labeled as a nnU-net modification. The learning rate in a deep learning network controls how quickly the model changes to incorporate the new patterns being presented. A rate that is too low will progress very slowly, however; a rate that is too high can cause unstable behavior. In addition to testing an adaptive learning rate, we also tested using a lower learning rate than was used in the original training.

Finally, nnU-net provides code for utilizing overlapping sliding windows during inference. The overlapped windows predictions are each weighed by a Gaussian matrix the same size as the window. The Gaussian matrix has higher weights at the center of the matrix and lower weights at the edges. The reason that using these overlapping windows should increase the performance is that predictions at the edges of volumes might be less accurate due to edges having less surrounding information then the center of volumes and thus should be weighted less.

An additional parameter we tested was fine-tuning only some layers of the model. When transfer learning with a smaller dataset, it is often best practice to fine-tune only some layers of the existing model to avoid overfitting [20]. Previous studies have investigated which layers of U-net should be trained for optimal performance, and found that training the encoding layers of a U-net model resulted in a higher dice score than training the decoding layers [21]. Since we are using a small training dataset, we tested training only the first, middle, and last two convolutional layers, as well as training all the U-net layers.

Our study endeavored to optimize transfer learning using principles from the nnU-net framework as well as other principles for fine-tuning U-net. We compared the performance in identifying axons in expert-annotated mouse brain slices both before and after fine-tuning on our annotated data. We also compared the performance of fine-tuning using the standard settings from the TrailMap package [15] against the TrailMap package with the preprocessing and post-processing modifications suggested by nnU-net [19, 22] and other U-net literature.

## Materials and Methods

### Animal Experiments

All animal experiments and handling were conducted in accordance with the US National Institutes of Health Guide for the Care and Use of Laboratory Animals and were approved by the Institutional Animal Care and Use Committee (IACUC) of Oregon Health & Science University (#IP00000955). All genetic reagents were used according to the protocols approved by Institutional Biosafety Committee (IBC) (# IBC-10-40). DbhCre mice were generously gifted by Dr. Patricia Jenson at NIHES (also available from The Jackson Laboratory; stock #033951) [23] and maintained by backcrossing with wildtype *C57Bl/6* mice acquired from Charles River, or home-bred within 5 generations of Charles River breeders. Mice 1-3 months old of both sexes with approximately equal proportions were used and were maintained on a 12-hour light-dark cycle, with food and water available ad libitum.

### Stereotaxic viral injections

Injections were performed using our established procedures [24]. Briefly, *Dbh^Cre+/-^* mice at 6 weeks of age were administered a pre-operative analgesic agent (carprofen; 5 mg/kg, SC), anesthetized with isoflurane (4% induction, 1-2% maintenance), and stabilized in a custom stereotaxic apparatus with body temperature being maintained by a heating pad. A small craniotomy was made over each target of interest and a pulled glass micropipette (Drummond, tip diameter: 15-20 µm), beveled sharp and loaded with the injectant, was lowered into the brain. Pipettes were front-filled with AAV (serotype 2/1) expressing either CAG-FLEX-GFP (UPenn, lot# V0827) or pCAG-FLEX-tdTomato-WPRE (Addgene cat# 51503). 3 min after reaching the target, 30-50 nL of virus was dispensed using a hydraulic injector (Narishige) followed by a 5-min waiting period. The pipette was retracted 0.3 mm, paused for 10 min, and then fully retracted. Up to two injections were made per mouse targeting the locus coeruleus bilaterally (in mm relative to Bregma; 4.80 posterior, +/- 0.75 lateral, 4.00 ventral). Following surgery, mice were given supplemental heat and monitored until ambulatory (typically 10-30 min) before being returned to their home cage. Additional daily doses of carprofen (5 mg/kg, SC) were given for the first 3 post-operative days. Mice were monitored daily for signs of infection, swelling, discharge, or changes in weight—if any clinical signs of concern became present (e.g., >20% weight loss within 3 days; body condition score <3/5; prolonged hunching, distention, lethargy, or diarrhea; tremors, spasticity, or seizures; frank bleeding; or other indications of moribund condition) mice would be immediately and humanely euthanized and excluded from further study. All animals injected for this study (n = 7) reached the planned experimental endpoint (see below) without adverse incident.

### Tissue collection

Three (3) weeks after viral infection, mice were deeply anesthetized with isoflurane and perfused transcardially with 50 mL room-temperature (RT) phosphate-buffered saline (PBS) followed by 50 mL of ice-cold 4% paraformaldehyde (PFA, w/v) and an additional 30 mL of RT PBS. The extracted brain was post-fixed in 4% PFA at 4 °C overnight under gentle agitation, rinsed 3x in RT PBS for 1 h under agitation, and then finally stored in PBS with 0.02% NaN_3_ at 4 °C.

### Tissue dilapidation, labeling, and clearing

Fixed brains were processed using a recent optimization of the AdipoClear framework (protocol v1.0 from https://mab3d-atlas.com) [25, 26]. All steps used gentle agitation at RT unless stated otherwise. Samples are first washed 3x (2 h, 4 h, overnight) in B1n buffer (in H_2_O: 0.1% Triton X-100, 2% glycine, 0.01% 10N NaOH, 20% NaN_3_) to block excess PFA and preserve antigens. Next, samples were delipidated with SBiP buffer (200µM Na_2_HPO_4_, 0.08% sodium dodecyl sulfate, 16% 2-methyl-2-butanol, 8% 2-propanol in H_2_O (pH 7.4)) for 1 h, 2 h, 4 h, overnight, and 3x 1 d. Samples were then rehydrated overnight in B1n buffer, then rinsed 4x (1 h, 2 h, 4 h, overnight) in PBS w/ NaN_3_. To begin immunolabeling, brains were first blocked with PTxwH buffer (in PBS: 0.1% Triton X-100, 0.05% Tween-20, 0.002% w/v heparin, 20% NaN_3_) for 4x (1 h, 2 h, 4 h, overnight). Samples were then incubated with primary antibodies (rabbit polyclonal anti-RFP, Rockland cat# 601-401-379, lot# 42872, 1:500; goat polyclonal anti-GFP, Rockland cat# 600-101-215, lot# 35577, 1:500) diluted in PTxwH for 20 d at 37 °C, then rinsed 5x (1 h, 2 h, 4 h, 12 h, 24 h) in PTxwH at RT. Next, samples were incubated with secondary antibodies (donkey polyclonal anti-rabbit Alexa 594, Jackson Immunoresearch cat# 711-587-003, lot# 148972, 1:150; donkey polyclonal anti-goat Alexa 647, Jackson Immunoresearch cat# 705-607-003, lot# 150295, 1:150) diluted in PTxwH for 18 d, and rinsed 5x (1 h, 2 h, 4 h, 12 h, 24 h) in PTxwH. Samples were then bleached in 0.3% H_2_O_2_ at 4 °C overnight, and washed 3x in 20 mM PB (16 mM Na_2_HPO_4,_ 4 mM NaH_2_PO_4_ in H_2_O) at RT for 2 h. For further delipidation, samples were dehydrated in an ascending gradient (20%, 40%, 60%, 80%) of MeOH in H_2_O for 1 h each, 3x (1 hours, 1.5 hours, 2 hours) 100% MeOH, then a descending gradient (80%, 60%, 40%, 20%) of MeOH in H_2_O for 1 h each. Next, brains were washed twice (2 h, 4 h) in 20 mM PB and finally twice in PTS solution (25% 2,2’-thiodiethanol/10 mM PB) (2 h, overnight), then equilibrated with 75% histodenz buffer (Cosmo Bio USA AXS-1002424) with refractive index adjusted to 1.53 using 2,2’-thiodiethanol. Samples were stored at ∼20 °C until acquisition.

### Light-sheet microscopy

The cleared brain samples were imaged horizontally with tiling using the LifeCanvas SmartSPIM light-sheet microscope. 561/647 nm lasers were used for Alexa 594/647 imaging with the 3.6×/0.2 NA detection lens. Light-sheet illumination is focused with NA 0.2 lenses from each side, and axially scanned with an electrically tunable lens coupled to the sCMOS camera (Hamamatsu Orca Back-Thin Fusion) in slit mode. The camera exposure was set at fast mode (2 ms) with 16-bit imaging. Image stacks were acquired along the dorsoventral axis following a 4 x 6 grid configuration. The x/y sampling rate was 1.866 μm and z step at 2 μm. Individual images from each stack originating from the same z-plane were stitched together using the TeraStitcher module (https://abria.github.io/TeraStitcher/) [27] resulting in ∼3600 horizontal image planes spanning the whole brain.

### TrailMap Imaging

The image labeled cube subsets used in this study are 160 pixels in the x, y, and z direction (∼ 320 x 320 x 300 µm^2^), with corresponding 188 pixel^3^ padded source image input cubes. All cubes originated from the same set of horizontal sections (∼4.5 mm below dorsal origin) and were selected at x/y positions that captured differing densities or “textures” of axonal labeling for training (See Figure 3 in the results section). For each cube, every 20th section is annotated manually, starting at the 15th slice.

**Figure 3.**
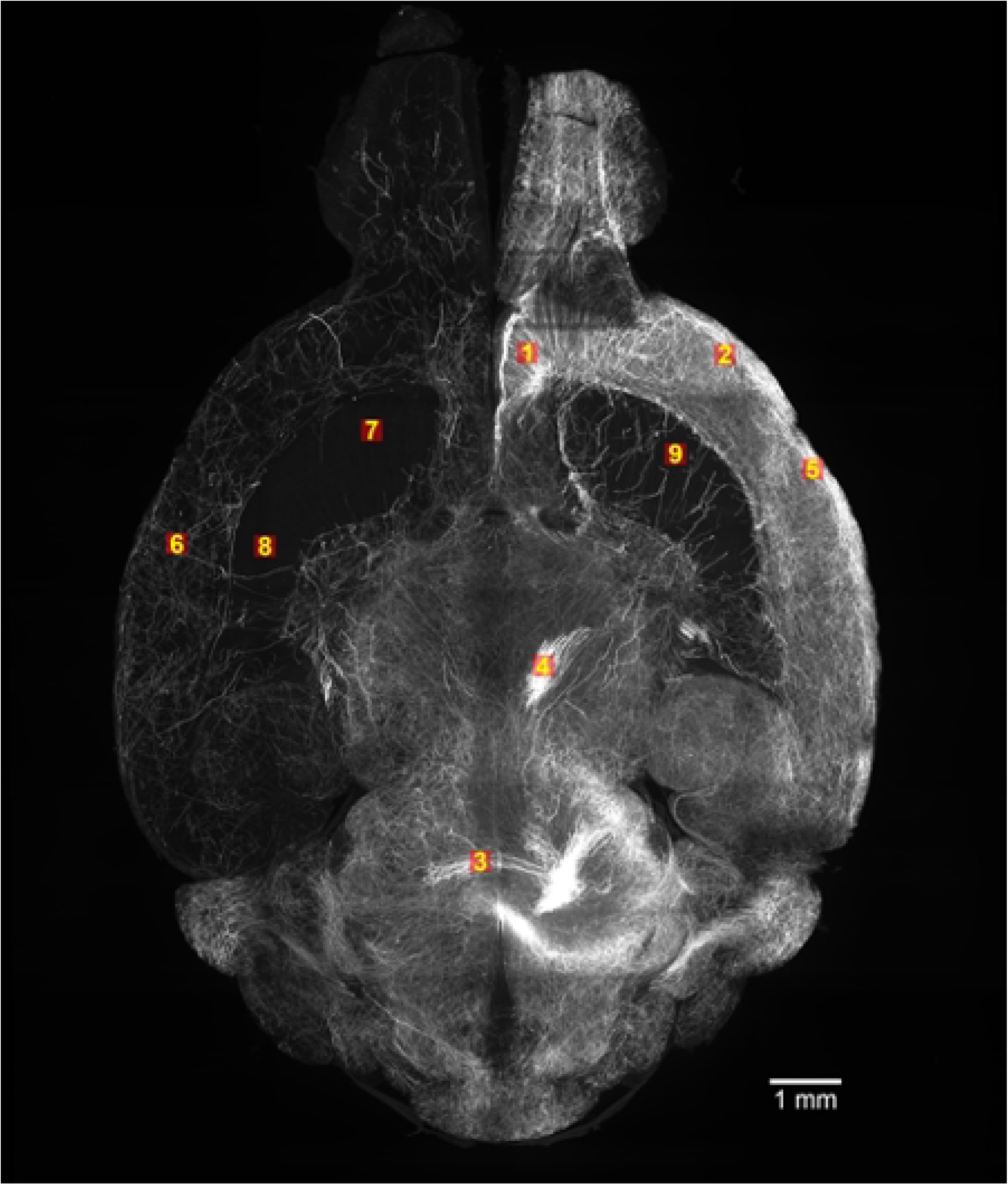
Location of labeled data. Nine 160 pixel^3^ labeled cubes were available in total. Seven cubes were used for training, one cube for validation, and one cube for testing. In the first experiment, cubes 3- 9 were assigned to training, cube 2 to validation, and cube 1 for testing. We also trained a second model with cubes 2- 4, 6-9 for training, cube 1 for validation, and cube 5 for testing.

For expert annotation, in Fiji, an image processing package, we enhanced the contrast with these settings: saturated: 0.35%, normalized: true, equalize: false, process all slices: true, use stack histogram: true, and exported the stack for loading in TrakEM2 [28]. In TrakEM2, a modeling program, we applied the CLAHE filter with settings: blockRadius: 63, bins: 255, slope: 3.0, and fast: true [29]. Sections were annotated using the AreaList interface and exported as binary label images.

After the expert annotation, the label images were processed with Trailmap’s axon edge annotator script to provide the edge labels. These images were subsequently used for training, validation, and testing of the modified TrailMap model.

### TrailMap Fine-tuning

We used the original TrailMap codebase from GitHub (https://github.com/AlbertPun/TRAILMAP) with our modifications [15]. We loaded the original pretrained model from TrailMap and fine-tuned it with our dataset using 100 epochs with batch-size of 6. Nine 160 pixel^3^ labeled cubes were available in total, with corresponding 188 pixel^3^ input cubes (Figure 3). Seven cubes were used for training, one cube for validation, and one cube for testing. In the first experiment, cubes 3-9 were assigned to training, cube 2 to validation, and cube 1 for testing. We also trained a second model with cubes 2- 4, 6-9 for training, cube 1 for validation, and cube 5 for testing. Our rationale for these assignments are as follows. Cubes 7-9 had no labeled axons and none of the cubes had labeled artifacts, so they were assigned to the training set in both experiments. Cubes 1, 2, and 5 are densely populated, so we intentionally randomly used one of each in the training, validation and testing data so the training dataset would include data similar to the validation and test dataset. The training data also included one sparse populated cube (cube 6) and two mediumly populated cubes (cubes 3 and 4). We used 100 3D volume samples from each cube, each with a 64-pixel^3^ input dimension and 32-pixel^3^ output dimension. The smaller size of the output is a feature of the TrailMap code – the model predicts axons only for the centers of the input volumes to make use of pixels around of the predicted area. The volumes are generated by a TensorFlow generator during the training.

For each epoch, the number of validation steps was the number of volumes (700 for training, 100 for validation) divided by the step size (6). We used an Adam optimizer and cross-entropy as the loss function, as used in the original TrailMap training scheme. The loss function weighted the classes for the axons, backgrounds, artifacts, and edges as 1.5, 0.2, 0.8, and 0.05, respectively. The model was saved during training at the highest weighted validation cross-entropy.

### Data Augmentation

The nnU-Net augmentations were implemented by incorporating the “augment_spatial” function from Batchgenerators [30]. TrailMap includes code for rotations and horizontal, vertical, and depth-wise flipping of volumes and rotations. Due to ease of use with elastic deformation, we used the Batchgenerators code for both rotations and elastic deformation, but used the TrailMap code for flipping.

We continued using the TrailMap method of intensity augmentation. The intensity was divided by a maximum intensity value and the training data was randomly scaled and summated with a random constant.

### Foreground Over-Sampling

The nnU-net code for oversampling randomly over-samples from the “foreground” data instead of all the data [19]. The foreground data in TrailMap are labeled as axons.

Within nnU-net, if a randomly selected volume extends past the limits of the cube, the volume is padded. With our input cubes sized at 188-pixel^3^ and volumes sized at 64- pixel^3^, this method would result in a high percentage of padded data. Instead, we continued to use the TrailMap sampling method, where the top, left, back corner of the training volume is selected randomly from the full image to create the training volumes. The top-left-back corner is selected from an area at least one training-volume dimension (64 pixels) from the opposite edges to ensure the cropping will produce a 64- pixel^3^ training volume. When oversampling axons, we selected the top, left, back corner pixels from a list of axon-labeled pixels 30% of the time.

### Learning rate

nnU-net uses a learning rate with a polynomial decay with 1000 epochs and a starting learning rate of 0.01. Since we are fine-tuning rather than training, and only are using 100 epochs and a lower starting learn rate, we tested reducing the learning rate by a factor of 0.1 only after a metric has stopped improving with a built-in TensorFlow learning-rate scheduler, ReduceLROnPlateau [31] as a nnU-net modification. TrailMap had a learning rate of 1e-3, but we also tested a lower learning rate of 1e-4

### Trainable Layers

We tested fine-tuning only the first, middle and last two 3D convolutional layers, as well as training with all the layers. When fine-tuning two layers, we did not freeze the batch-normalization layer.

### Gaussian Weighted Overlapping Windows

Our modified TrailMap code implemented the same logic and the Gaussian matrix function as nnU-net to calculate the value of overlapping windows. However, we did not extend the Gaussian matrix function with a Gaussian prediction for mirrored images as implemented in nnU-net [19]. In the Gaussian matrix function, each overlapping volume is multiplied by a Gaussian matrix before being added to the predicted image. Each resulting predicted value is then divided by the sum of the Gaussian multipliers used at that location in the image. The final value at each voxel by a weighted sum, with the weights representing the centrality of the voxel of each volume.

### Performance Metrics

For the performance metrics in this paper, the TrailMap inference function was used to predict axons for the validation and test data cubes. The TrailMap inference function segments the image in a sliding window. The input image used by TrailMap is larger than the segmented output (64 pixel^3^ input vs 36 pixel^3^ output), so the segmented image is smaller than the input image by an offset at the edges (64 pixel^3^ input – 36 pixel^3^ output/2 = 14-pixel offset). Therefore, we used a 188 pixel^3^ input to predict a 160 pixel^3^ output. The performance metrics were obtained from the labeled input and the segmented output for both the validation and test datasets.

The metrics used were:

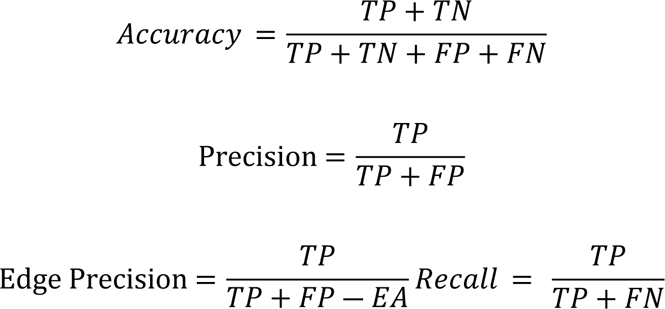

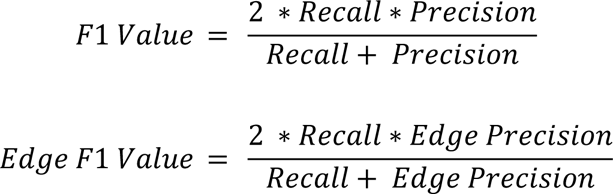

with Edges identified as Axons (EA), True Positive (TP), True Negative (TN), False Positive (FP), and False Negative (FN).

The authors of TrailMap noted that they did not include edges when calculating performance metrics [15]. To remain consistent with the original code, we also calculated “Edge Precision” and “Edge F1-score”, where the edges classified as axons were not included in the false positives. We used Edge F1-score as our primary performance evaluator so both recall and precision would be considered. Using just the recall would weigh the False Negatives while disregarding any False Positives, and using only precision would weigh the False Positives while disregarding any False Negatives.

## Results

After training, we segmented the validation datasets to determine the optimal modifications to the TrailMap code. We only used modifications that lead to a stable lower validation loss than the default model. After selecting the best modifications, we compared the performance on the test dataset metrics of both the best fine-tuned TrailMap model and the original model.

### Freezing layers

The performance metrics of the model after fine-tuning all the layers or the first, middle, and last two convolutional layers of the neural network on the validation dataset are shown in Table 1. Figure 4 shows the validation loss over 100 epochs. The model with the fine-tuned last layers lead to the lowest, stable validation loss and lead to a higher Edge F1-score of 79.6% compared to 78.4% for the next best model.

**Figure 4.**
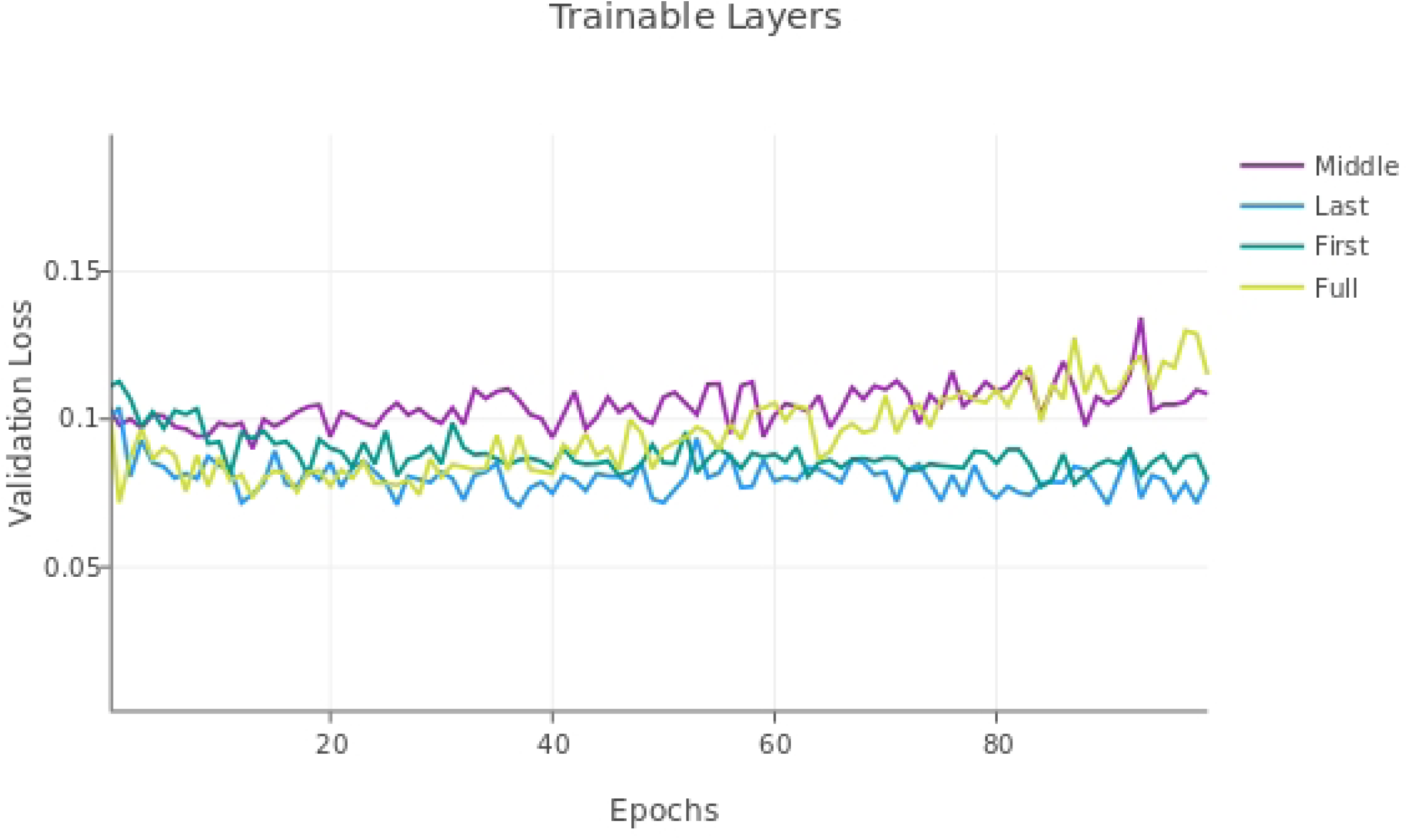
Validation loss with fine-tuning all the layers (Full) or the first, middle, and last two convolutional layers of the neural network. All models had a learning rate of 1e-4.

**Table 1.** Performance metrics with 1) all U-net layers trainable (Full) 2) the last two layers (decoder) trainable (Last), 3) the middle two layers (Middle), and the first two layers (First) with the validation dataset. All models had a learning rate of 1e-4. Original indicates the results of the original model without fine-tuning. Models were trained with cubes 3-9, validated with cube 2, and tested with cube 1.

| Training Layers | Accuracy | Precision | Edge Precision | Recall | F1 Score | Edge F1 Score |
| --- | --- | --- | --- | --- | --- | --- |
| Original | 86.5% | 52.6% | 70.5% | 85.5% | 65.1% | **77.3%** |
| Full | 84.9% | 49.2% | 68.6% | 91.5% | 64.0% | **78.4%** |
| Last | 84.5% | 48.7% | 69.2% | 93.7% | 64.1% | **79.6%** |
| Middle | 83.9% | 47.6% | 65.5% | 90.6% | 62.4% | **76.0%** |
| First | 84.0% | 47.8% | 66.9% | 92.6% | 63.1% | **77.7%** |

### Modifying loss rate

The performance metrics of the model with a learning rate of 1e-3 or 1e-4 and with the last two layers fine-tuned are shown in Table 2 for the validation dataset. Figure 5 shows the validation loss over 100 epochs. The model with the 1e-4 learning rate had the lowest stable validation loss value, so, although the performance metrics of the adjusted F1-score was slightly higher for 1e-4 vs the 1e-3 learning rate (80.2% compared with 79.6%), we continued to use a 1e-4 learning rate.

**Figure 5.**
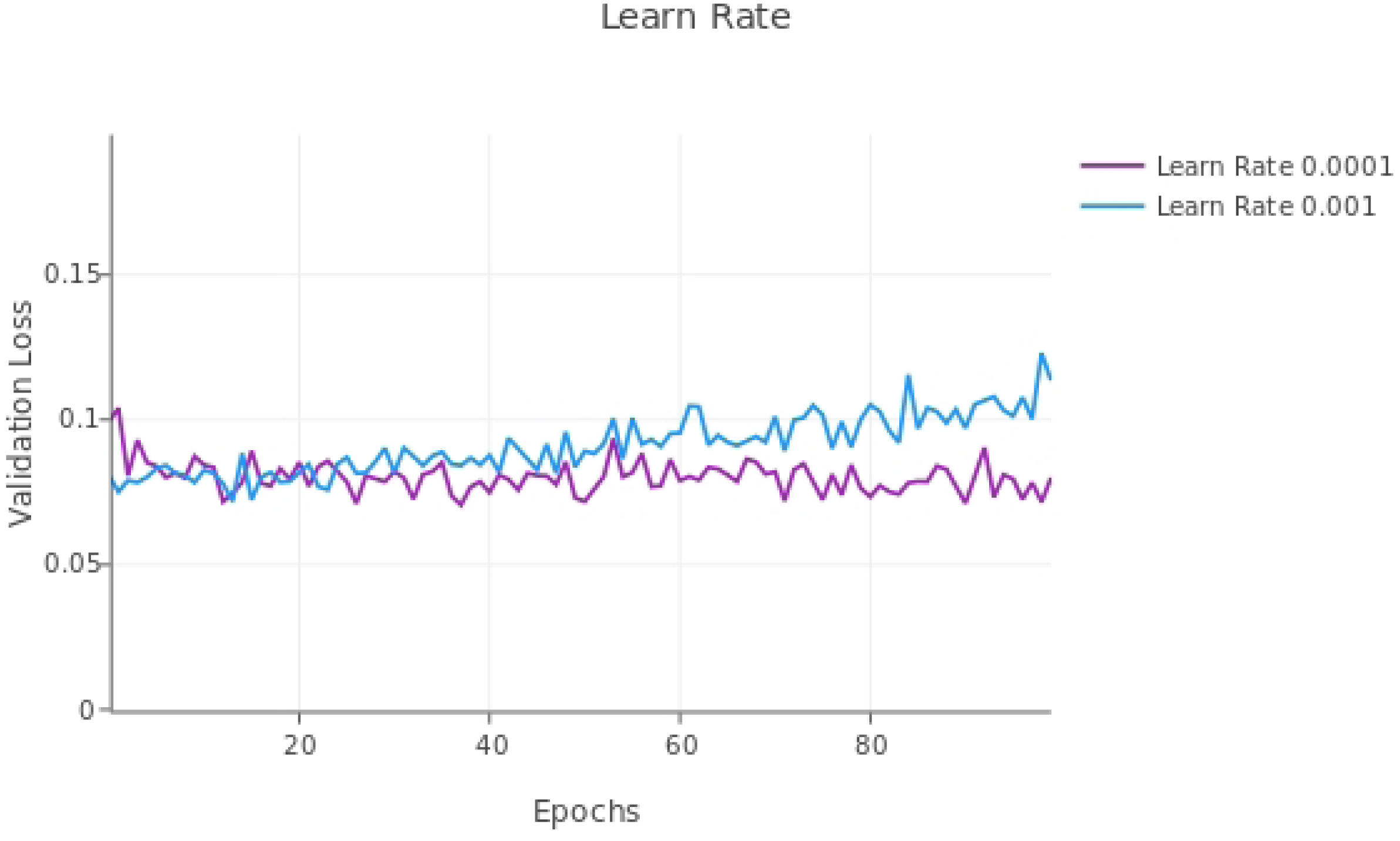
Validation loss with fine-tuning the last two layers of the TrailMap model with different (non-adaptive) learning rates.

**Table 2.** Performance metrics with the 1e-4 or 1e-3 learning rate using the validation dataset cube 2. All models evaluated below only had the last two layers fine-tuned. “Original” in the first column below indicates the results of the original TrailMap model without fine-tuning. Models were trained with cubes 3-9, validated with cube 2, and tested with cube 1.

| Learn rate | Accuracy | Precision | Edge Precision | Recall | F1 Score | Edge F1 Score |
| --- | --- | --- | --- | --- | --- | --- |
| Original | 86.5% | 52.6% | 70.5% | 85.5% | 65.1% | **77.3%** |
| 1e-4 | 84.5% | 48.7% | 69.2% | 93.7% | 64.1% | **79.6%** |
| 1e-3 | 85.6% | 50.6% | 70.7% | 92.7% | 65.5% | **80.2%** |

### nnU-net Modifications Models

The results for single modifications are shown in Table 3. The nnU-net modifications used were: over-sampling axons, using an adaptive learning rate, elastic deformation, and rotations, in addition to the built-in TrailMap augmentation of flipping the volumes. The loss curve shows no modification resulted in a consistently lower validation loss curve, both for the augmentation modifications (Figure 6) and the other modifications (Figure 7).

**Figure 6.**
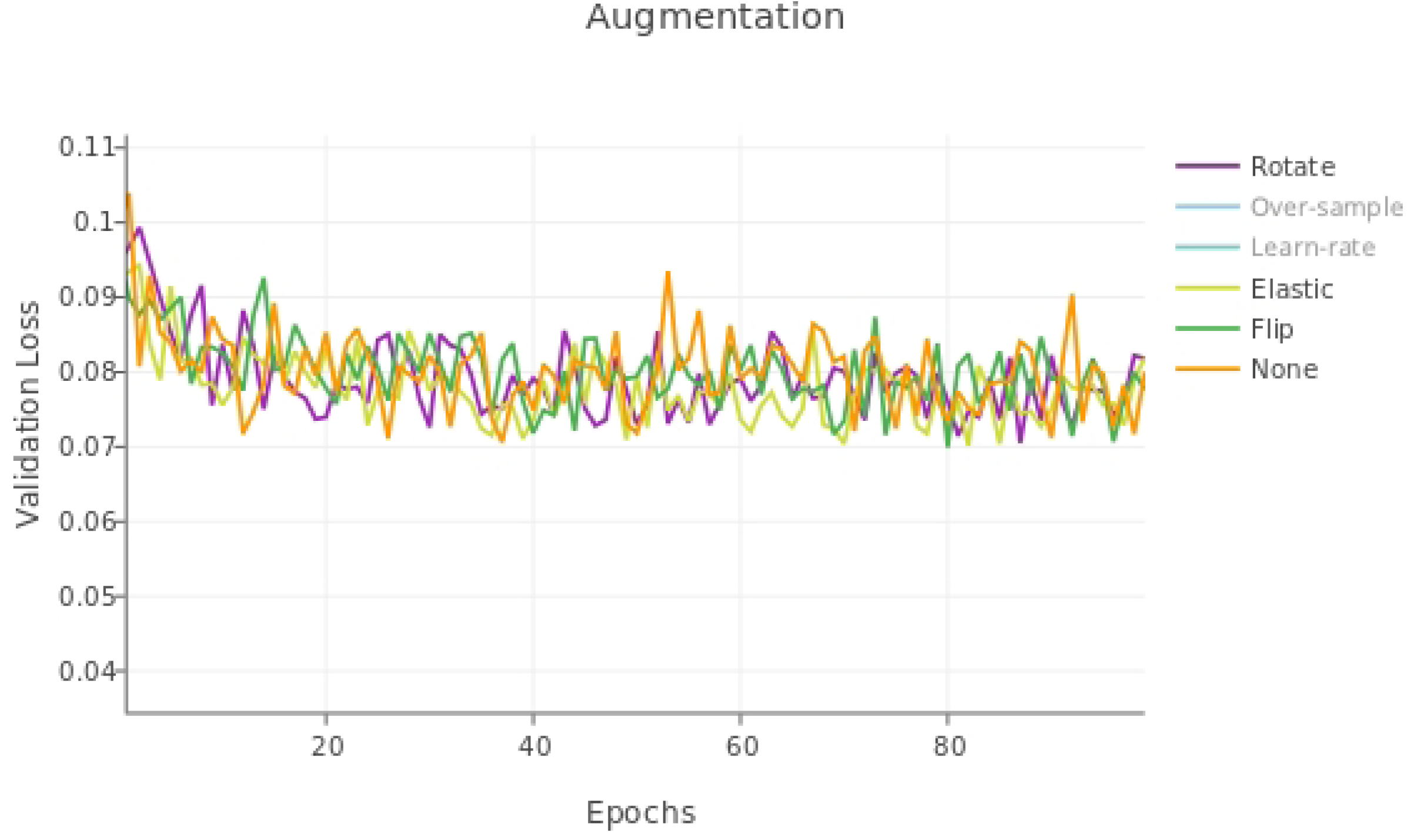
Validation loss for single nnU-net augmentations.

**Figure 7.**
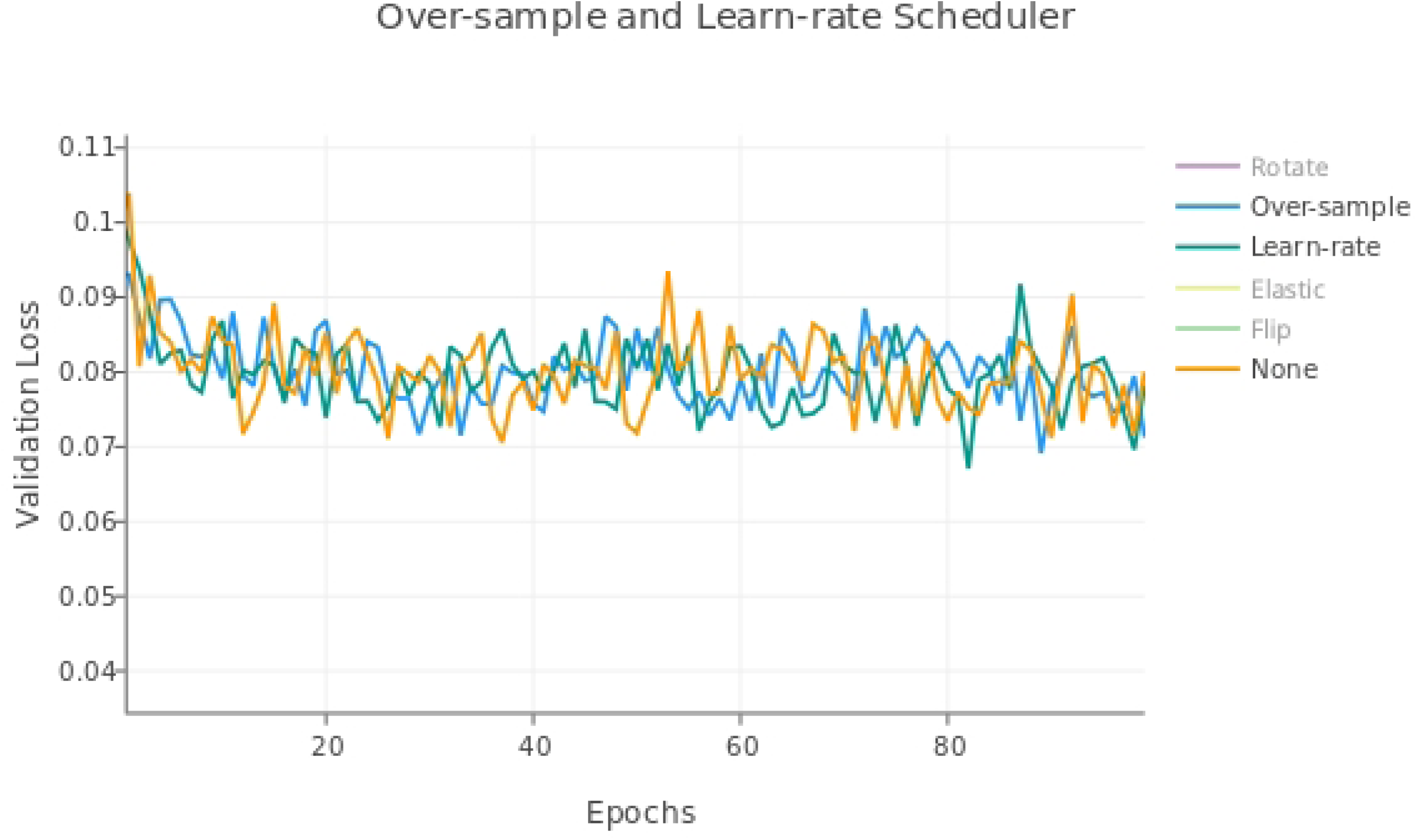
Validation loss for single nnU-net modifications, excluding augmentations.

**Table 3.**
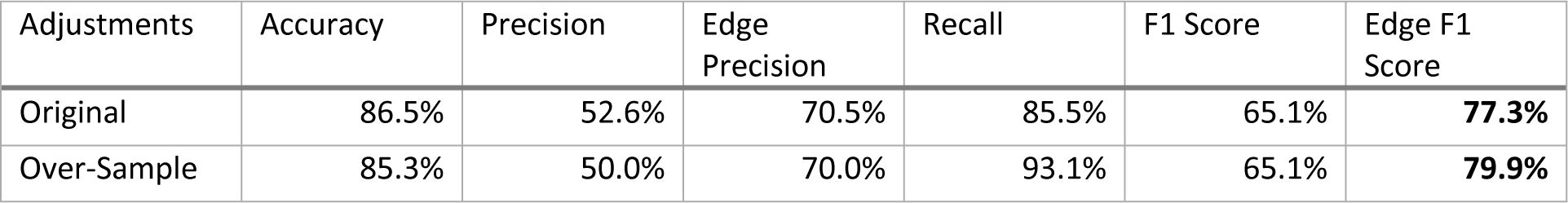
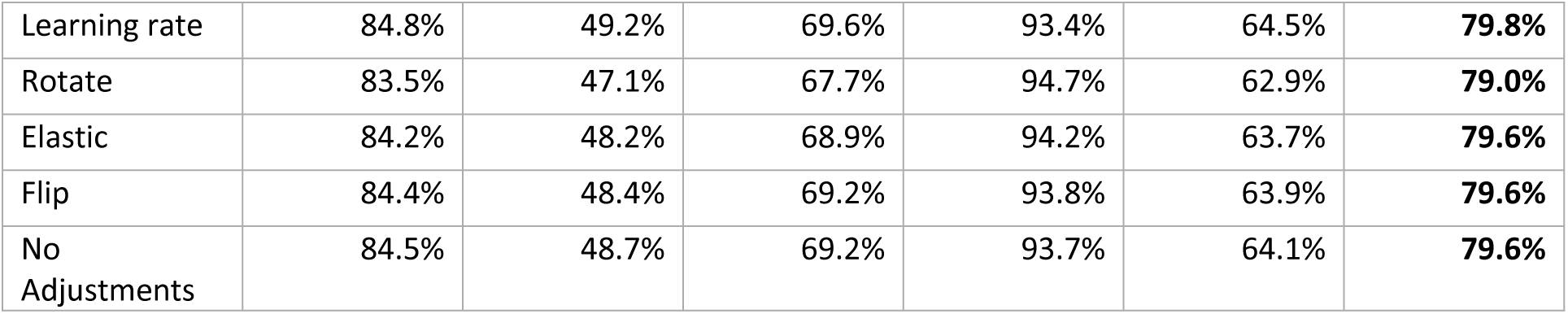
Performance metrics for models with over-sampling, learn rate scheduler, rotation, elastic deformation, and flipping for the validation dataset. All models below only had the last two layers fine-tuned with a learning rate of 1e-4 (or starting learning rate in the case of the adaptive learning rate). Original indicates the results of the original model without fine-tuning. Models were trained with cubes 3-9, validated with cube 2, and tested with cube 1.

| Adjustments | Accuracy | Precision | Edge Precision | Recall | F1 Score | Edge F1 Score |
| --- | --- | --- | --- | --- | --- | --- |
| Original | 86.5% | 52.6% | 70.5% | 85.5% | 65.1% | **77.3%** |
| Over-Sample | 85.3% | 50.0% | 70.0% | 93.1% | 65.1% | **79.9%** |
| Learning rate | 84.8% | 49.2% | 69.6% | 93.4% | 64.5% | **79.8%** |
| Rotate | 83.5% | 47.1% | 67.7% | 94.7% | 62.9% | **79.0%** |
| Elastic | 84.2% | 48.2% | 68.9% | 94.2% | 63.7% | **79.6%** |
| Flip | 84.4% | 48.4% | 69.2% | 93.8% | 63.9% | **79.6%** |
| No Adjustments | 84.5% | 48.7% | 69.2% | 93.7% | 64.1% | **79.6%** |

### Best Overall Model

We compared the performance of the best fine-tuned TrailMap model with the original model on the held-out test set. The best model only had the last two layers fine-tuned with a learning rate of 1e-4, and none of the nnU-net modifications. Table 4 shows that the best fine-tuned TrailMap model had a higher Edge F1-score than the original model (91.6% vs 86.7%). Figure 8 shows the color-coded true positives, false positive, and true negatives of the predictions of each model, with and without Gaussian overlap. It’s visible in Figure 8 that the original model has more false negatives then the fine-tuned model, while the fine-tuned model has more false positives.

**Figure 8.**
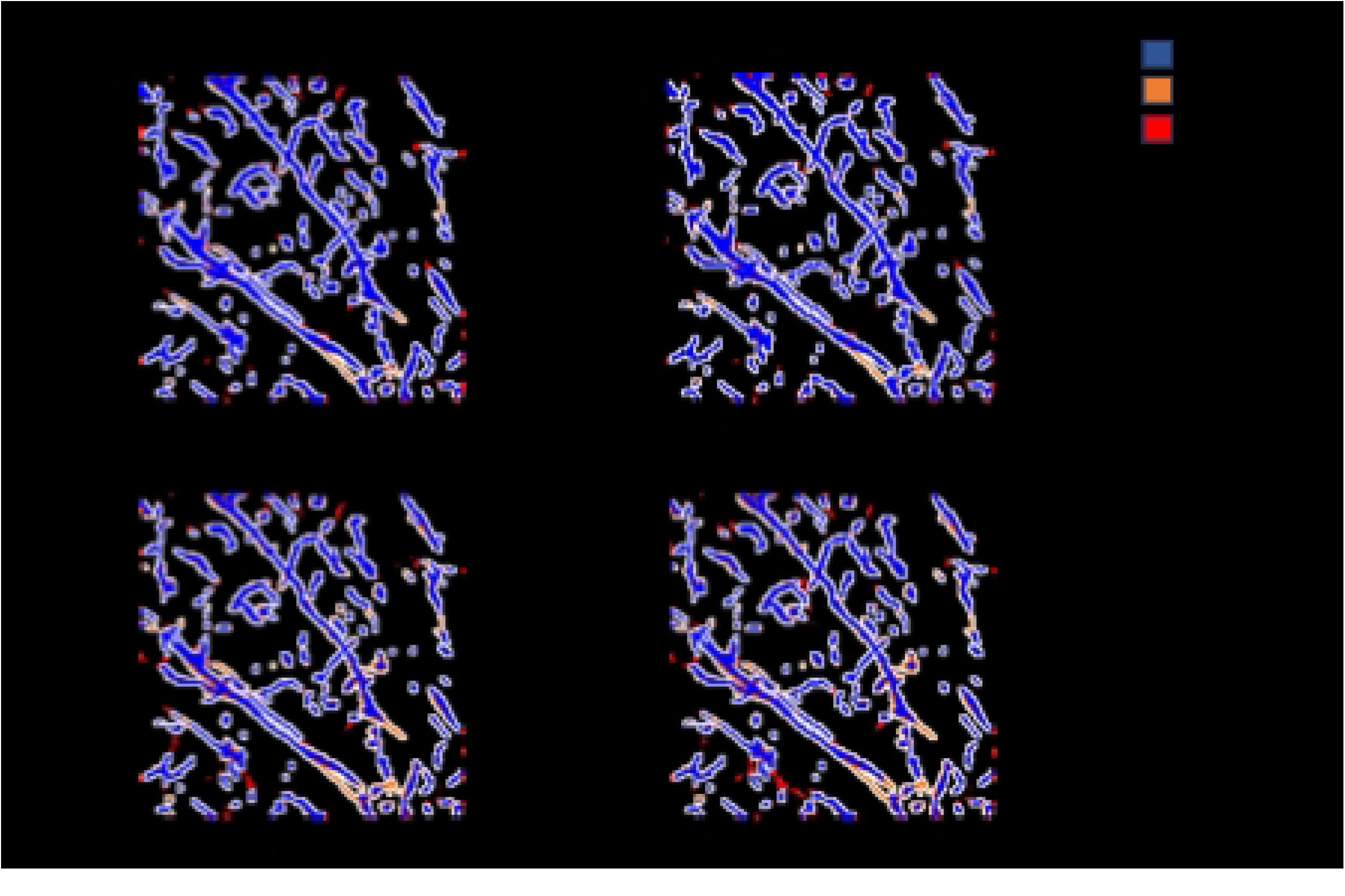
Cube 1 test dataset slice predictions created with A) best fine-tuned model B) best fine-tuned model with Gaussian overlap inference, C) best original model D) best original model with Gaussian overlap inference. The dimensions are 298.56 um on x and y axis.

**Table 4.** Performance metrics for the best model, which is trained with only the last two layers fine-tuned and a learning rate of 1e-4 but with no further adjustments, on the test dataset compared with the original model without fine-tuning. Models were trained with cubes 3-9, validated with cube 2, and tested with cube 1.

| Training Layers | Learn rate | Accuracy | Precision | Edge Precision | Recall | F1 Score | Edge F1 Score |
| --- | --- | --- | --- | --- | --- | --- | --- |
| **Original** |  | 92.5% | 68.6% | 92.0% | 81.9% | 74.7% | **86.7%** |
| **Last** | 1e-4 | 92.8% | 67.8% | 93.9% | 89.4% | 77.1% | **91.6%** |

A visualization of the predictions of the best and original TrailMap model (Figure 8B and 8C) is qualitatively similar to the input image (Figure 8A).

We trained and validated the original model with cube 1 as the validation dataset and cube 5 as the test dataset, the adjusted F1-score was slightly higher for the original model than for the fine-tuned (Table 5) (84.8% vs 84.4%). Consistent with the previously trained model, the recall was higher, and the precision was lower for the fine-tuned model compared with the original model.

**Table 5.** Performance metrics for a new model, trained with on datasets 2-4, 6-9, validated with cube 1, and tested with cube 5. The metrics for the validation and test dataset are shown for the original model and the model trained with the previously determined best parameters.

| Data | Training Layers | Learn rate | Accuracy | Precision | Edge Precision | Recall | F1 Score | Edge F1 Score |
| --- | --- | --- | --- | --- | --- | --- | --- | --- |
| **Val** | Original |  | 92.5% | 68.6% | 92.0% | 81.9% | 74.7% | **86.7%** |
| **Val** | Last | 1e-4 | 92.6% | 66.8% | 93.9% | 90.2% | 76.7% | **92.0%** |
| **Test** | Original |  | 84.9% | 49.8% | 81.4% | 88.4% | 63.7% | **84.8%** |
| **Test** | Last | 1e-4 | 80.6% | 43.4% | 75.6% | 95.4% | 59.7% | **84.4%** |

### Gaussian-weighed Overlap

Table 6 indicates if using Gaussian-weighed overlapping windows increases or decreases the accuracy or Edge F1-score. Using a Gaussian-weighed overlap for inference, where the Gaussian matrix weights each voxel by the centrality of the voxel within the current window, generally increased the model accuracy and adjusted F1-score. Figure 8 shows inference created with Gaussian-weighted overlapping windows in panels C and D. In Table S1, we compared Gaussian inference with the default inference metrics for models that were not run with the test dataset in Table 4 and found that although the accuracy was consistently improved with Gaussian inference, the Edge F1 score often decreased.

**Figure 9.**
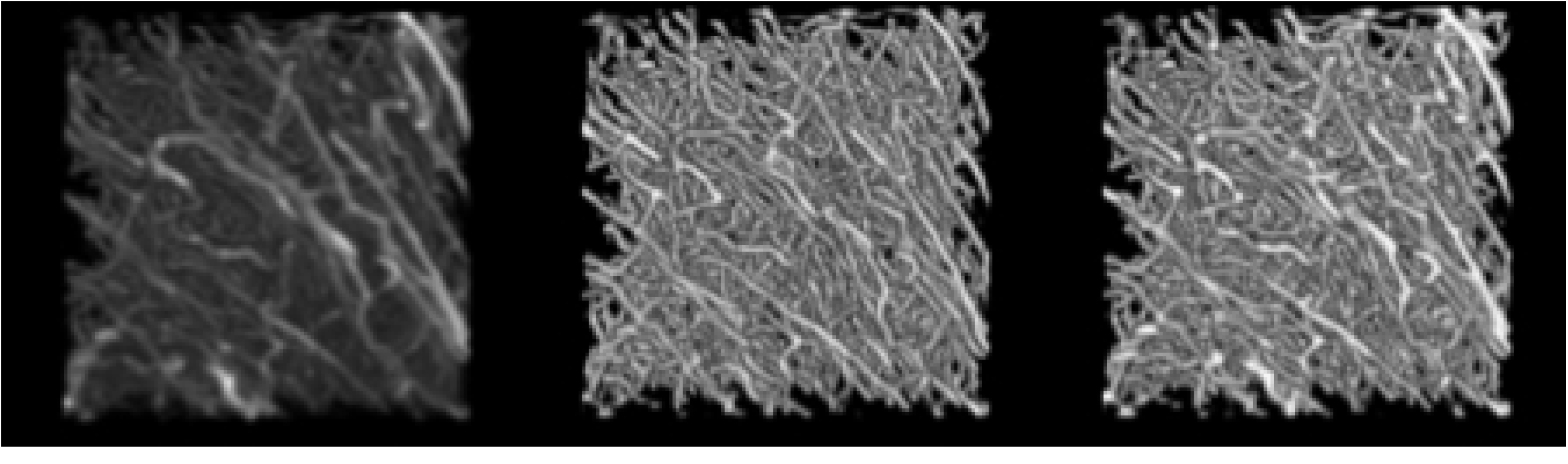
Data Inference results where A) is the input image, B) is the segmentation image result produced using the original model, and C) is the segmentation image result produced using the best fine-tuned model for cube 1 test dataset. The input image was processed by the CLAHE filter as described in the methods. Only segmentation values above 0.5 are shown. Displayed with Fiji. The dimensions are 298.56 um on x and y axis and 320 um on the z axis

**Table 6.** Effect of using Gaussian (G) on all results as measured by accuracy and Edge F1-scores. The difference column indicates if the model used 1) one of the nnU-net modifications (oversample, rotate, elastic, an adaptive learning rate, flip, and all modifications) 2) a higher learning rate 3) different training layers, 4) the original model or the model with the best parameters, or 5) the original or model with best parameters trained, tested, and validated with different cubes (with cube 1 as the validation dataset and cube 5 as the test dataset). The model with the best parameters had none of the nnU-net modifications, but was trained with the last two layers and had a learn rate of 1e-4. Imp: Improvement

| Data | Difference | Layers | LR | Accuracy | G | Imp | Edge F1 | G | Imp |
| --- | --- | --- | --- | --- | --- | --- | --- | --- | --- |
| Val | Original |  |  | 86.5% | 86.8% | Yes | 77.3% | 77.7% | Yes |
| Val | Over-Sample | Last | 1e-4 | 85.3% | 85.4% | Yes | 79.9% | 80.0% | Yes |
| Val | Rotate | Last | 1e-4 | 83.5% | 83.7% | Yes | 79.0% | 79.0% | Yes |
| Val | Elastic | Last | 1e-4 | 84.2% | 84.4% | Yes | 79.6% | 79.6% | Yes |
| Val | Learning rate | Last | 1e-4 | 84.8% | 85.0% | Yes | 79.8% | 79.9% | Yes |
| Val | Flip | Last | 1e-4 | 84.4% | 84.5% | Yes | 79.6% | 79.7% | Yes |
| Val | Higher LR | Last | 1e-3 | 85.6% | 85.8% | Yes | 80.2% | 80.4% | Yes |
| Val | Training Layer | Full | 1e-4 | 84.9% | 85.0% | Yes | 78.4% | 78.7% | Yes |
| Val | Best Model | Last | 1e-4 | 84.5% | 84.7% | Yes | 79.6% | 79.7% | Yes |
| Val | Training Layer | Middle | 1e-4 | 83.9% | 84.2% | Yes | 76.0% | 76.5% | Yes |
| Val | Training Layer | First | 1e-4 | 84.0% | 84.4% | Yes | 77.7% | 78.0% | Yes |
| Val (cube 1) | Original |  |  | 92.5% | 92.4% | No | 86.7% | 86.5% | No |
| Val (cube 1) | Best Model | Last | 1e-4 | 92.6% | 92.7% | Yes | 92.0% | 92.0% | No |
| Test | Original |  |  | 92.5% | 92.4% | No | 86.7% | 86.5% | No |
| Test | Best Model | Last | 1e-4 | 92.8% | 92.9% | Yes | 91.6% | 91.7% | Yes |
| Test (cube 5) | Original | Original |  | 84.9% | 85.0% | Yes | 84.8% | 85.2% | Yes |
| Test (cube 5) | Best Model | Last | 1e-4 | 80.6% | 81.0% | Yes | 84.4% | 84.9% | Yes |

## Discussion

Often, the most challenging aspect of fine-tuning a LSM segmentation model is creating the training data. The labeling process is arduous and requires expert skill to identify the axons. The methods explored here provide a broad roadmap for users to fully utilize and improve upon existing trained models, reducing the burden of manual data annotations. In addition to the modifications to the TrailMap code, we have provided a new dataset of expert-annotated noradrenergic axons. The utility of TrailMap in axon segmentation has generalized efficiently to our dataset both with the original TrailMap model and with the use of fine-tuning and additional modifications. Recently, others have shown success in training a model with nnU-net to segment axons without manual annotation showing state-of-the-art performance[32]. While this new, automated strategy significantly reduces manual input in axon segmentation, the model is still reliant on extensive training data or transfer learning techniques, such as those described in this paper, to develop new training modules for specific axon types or sample batches.

While we did not find a consistent increase in the adjusted F1-score with our best fine-tuned model compared to the original model, we did find a consistent increase in recall and decrease in precision. Whether this trade-off of precision for increased recall is useful depends on the task. Perhaps in a task where further processing would involve expert-confirmation of axons, recall would be more important as the expert can more easily remove incorrect axons than find new ones. On the other hand, in situations where the density of axons must be guaranteed to be above a threshold, precision would be more important. Friedman et al. had previously successfully fine-tuned the model with 17 training cubes, rather than the 7 that we used [15]. We ultimately show that it is possible to have a positive effect on the model performance in some metrics even with a small training dataset.

Training only the last two convolutional layers of the model and using a lower learning rate had a positive impact on model performance. Previous studies have found that training the encoding layers (the first layers) of the model had the most positive effect, so our results are inconsistent with the results in Amiri et al. (2019), which may be due to the different image types and dataset size used in their fine-tuning training set compared to ours. Fine-tuning the first two encoding layers did lead to the second lowest stable validation loss.

The nnU-net modifications did not have a clear effect on the Edge F1-score of the model. Some of the Edge F1-scores were slightly higher or lower, but the validation loss curve makes it clear that no modification improves the model enough to result in a stable, lower validation loss curve. Generally, the augmentation of the dataset should mimic the variability of the test dataset. It is possible that the small size of our testing dataset (one cube) does not contain enough variability to benefit from augmentation, and that we would see an effect of augmentation with a larger test dataset.

The learning rate scheduler would be expected to increase the stability of the validation loss. We did not see this clearly in the validation loss, perhaps because we already used a lower learning rate as a starting point.

Over-sampling did not have a positive effect. Two possible reasons are that our oversampling method restricts the possible locations of the corners of the volumes to one of labeled slices rather than all slices of data. Another is that in our dataset, we already have a high percentage of axons, so over-sampling might be unnecessary.

The nnU-net modification of including sliding inference windows with Gaussian weights did generally improve our results. All but one of the models used in this paper were improved by Gaussian overlap as measured by accuracy (the model that was not improved, the original model, was shown multiple times in Table 6). However, there was not a consistent improvement as measured for the Edge F1-score. Accuracy might be a better measurement for Gaussian overlap improvement rather than Edge F1-score. This is because the trained mode penalizes misclassified edges less than misclassified axons, whereas the Gaussian overlap should increase the number of true positive and negatives regardless of the segmentation class. A possible reason Gaussian overlap did not perform better in all scenarios is that TrailMap already ensures no voxel was located at the edge of the input when it is segmented because it segments a smaller output from a larger input.

A further area of study could be to compare the performance on models trained on specific brain areas vs generalized models. There was a large difference in results between the validations dataset (best model - Edge F1 score: 79.6%) and the test dataset (best model - Edge F1 score: 91.6%). These results suggest that even within one brain, the variation warrants further validation in individual sections. This variation indicates the problem of complete mapping of neuronal structures into a co-registered system will likely require many structure-specific solutions in the short-term. In addition to providing a new fine-tuned model, we also provided a roadmap to test similar fine-tuning modifications on specific structures.

## Acknowledgements

We thank Amelia Culp for assistance with animal injections, and Dr. Patricia Jenson for providing *Dbh^Cre^* mice. MTO, RMEW, SA, MO, LMB, BJWR were supported by the Laboratory Directed Research and Development at Pacific Northwest National Laboratory (PNNL), a Department of Energy facility operated by Battelle under contract DE-AC05-76RLO01830. WW, KB, and ZW were supported in part by a NIH/BRAIN Initiative Grant RF1MH128969. MAM and TM were supported by two NIH/BRAIN Initiative Grants R01NS104944, RF1MH120119 and NIH R01NS081071. This research is affiliated with the Pacific northwest bioMedical Innovation Co-laboratory (PMedIC) collaboration between OHSU and PNNL.

